# Diversity of tRNA Clusters in the Chloroviruses

**DOI:** 10.1101/2020.07.06.190819

**Authors:** Garry A. Duncan, David D. Dunigan, James L. Van Etten

## Abstract

Viruses rely on their host’s translation machinery for the synthesis of their own proteins. Problems belie viral translation when the host has a codon usage bias (CUB) that is different from an infecting virus with differences in the GC content between the host/virus genome. Here, we evaluate the hypothesis that chloroviruses adapted to host CUB by acquisition and selection of tRNAs that at least partially favor their own CUB. The genomes of 41 chloroviruses comprising three clades of three different algal hosts have been sequenced, assembled and annotated. All 41 viruses not only encode tRNAs, but their tRNA genes are located in clusters. One tRNA gene was common to all three clades of chloroviruses, while differences were observed between clades and even within clades. By comparing the codon usage of one chlorovirus algal host, whose genome has been sequenced and annotated (67% GC content), to that of two of its viruses (40% GC content), we found that the viruses were able to at least partially overcome the host’s CUB by encoding tRNAs that recognize AU-rich codons. In addition, 39/41 chloroviruses encode a putative lysidine synthase, which alters the anticodon of tRNA^met^ that normally recognizes AUG to recognize the codon AUA, a codon for isoleucine. This is advantageous to the viruses because the AU-rich codon AUA is 12-13 times more common in the chloroviruses than their host. Evidence is presented that supports the concept that chlorovirus tRNA clusters were acquired prior to events that separated them into the three clades.

**IMPORTANCE:** Chloroviruses are members of a group of giant viruses that infect freshwater green algae around the world. More than 40 chloroviruses have been sequenced and annotated. In order to propagate efficiently, chloroviruses with low GC content must overcome the high GC content and codon usage bias (CUB) of their hosts. We provide support for one mechanism by which viruses can overcome host CUB. Specifically, the chloroviruses examined herein encode tRNAs whose cognate codons are common in the viruses but not in the host. Virus-encoded tRNAs that recognize AU-rich codons enable more efficient protein synthesis, thus enhancing viral propagation. The tRNA genes are located in clusters and the original tRNA gene cluster was acquired by the most recent common ancestor of the four chlorovirus clades. Furthermore, we show some conservation among all clades, but also substantial variation between and within clades, demonstrating the dynamics of viral evolution.

## INTRODUCTION

Viruses rely on most or all of their host’s translation machinery to synthesize their proteins. A conflict for viruses occurs when they have a codon usage bias (CUB) different from their host. However, some viruses have genes that help adapt to the host’s CUB in favor of their own CUB by encoding tRNAs. These include large dsDNA viruses infecting eukaryotic organisms (see Morgado and Vicente, 2019 for an extensive list) and bacteriophages (Abe *et al*., 2014). Recently, tRNA-encoding genes have also been reported in small ssDNA and ssRNA viruses (Morgado and Vicente, 2019).

Among the group of tRNA-encoding viruses are large viruses that infect algae (Michely *et al.,* 2013; Pagarete *et al*., 2013; Santini *et al*., 2013; Derelle *et al.*, 2015), including the chloroviruses (family *Phycodnaviridae*) with a lytic life-style (Li *et al*., 1997; Nishida *et al*., 1999; Lee *et al.,* 2005. Fitzgerald *et al*., 2007, 2007a, 2007b; Jeanniard *et al*., 2013). The chloroviruses have genomes that are 290 to 370 kb in size and are predicted to encode up to 400 proteins (CDSs) and 16 tRNAs (Van Etten *et al.*, 2020). They infect certain chlorella-like green algae that live in a symbiotic relationship with protists and metazoans (referred to as zoochlorellae), forming the holobiont. There are four known clades of chloroviruses based on the host they infect: viruses that infect *Chlorella variabilis* NC64A (referred to as NC64A viruses), viruses that infect *Chlorella variabilis* Syngen 2-3 (referred to as Osy viruses), viruses that infect *Chlorella heliozoae* SAG 3.83 (referred to as SAG viruses), and viruses that infect *Micractinium conductrix* Pbi (referred to as Pbi viruses).

The genomes of more than 40 chloroviruses, representing 3 of the 4 clades, have been sequenced, assembled and annotated (Jeanniard *et al*., 2013 and references cited therein). The genome GC content of the NC64A viruses ranges from 40%-41%; the GC content of the Pbi viruses ranges from 44-47%; while the GC content of the SAG viruses ranges from 48-52% (Jeanniard *et al.,* 2013). Their hosts, *C. variabilis* NC64A (Blanc *et al*., 2010) and *M. conductrix* Pbi (Arriola *et al*., 2018), both have nuclear genome GC contents of 67%, and it is assumed that the *C. heliozoae* has a similar GC content.

Two NC64A chloroviruses, PBCV-1 and CVK2, also have the interesting property of clustered tRNA genes (Lee *et al*., 2005; Nishida *et al*. 1999). Clusters of tRNA genes are found in some bacteriophage (Morgado and Vicente, 2019), as well as in a small percentage of organisms from all three domains of life (Bermudez-Santana *et al*., 2010; Morgado and Vicente, 2018, 2019, 2019a). Clusters of tRNA-genes have also been reported in mitochondria and plastids (Abe *et al*., 2014; Fan *et al*., 2017; Freidrich *et al*., 2012).

This current *in silica* investigation had several purposes. i) To evaluate chlorovirus tRNA genes with respect to clustering. ii) To compare and contrast the tRNA genes within and among the three clades of chloroviruses. iii) To determine if there was a CUB in the algal host *C. variabilis* NC64A, and likewise, if there was a CUB among the chloroviruses. iv) To evaluate whether the chlorovirus clades acquired their tRNA gene clusters before or after their divergences from their most recent common ancestor (MRCA).

## MATERIALS AND METHODS

### Genomic sequence data

In total, the genomes of 41 sequenced, assembled (in some cases to draft genomes) and annotated chloroviruses that represent three of the four known chlorovirus clades were used in this study: 14 NC64A viruses, 13 SAG viruses, and 14 Pbi viruses (Jenniard *et al*., 2013 and references cited therein). Table S1 provides the accession numbers, as well as the source of the viruses. The genome of *C. variabilis* NC64A, which is the host of the NC64A viruses, has been sequenced and annotated (project accession number ADIC00000000; Blanc *et al.,* 2010). *M. conductrix*, the host of the Pbi virus, has also recently been sequenced and annotated (Arriola *et al.,* 2018), but its sequence was not included in this study.

### Identification and localization of the tRNA genes

A tRNA gene cluster was previously reported for chlorovirus PBCV-1 (Lee *et al*., 2005; Dunigan *et al.,* 2013), as well as CVK2 (Nishida *et al*., 1999). The other 40 chloroviruses were then examined to determine if they also had tRNA gene clusters, and their tRNA genes and gene order were documented. The tRNA genes for each virus were identified by entering the genome accession numbers into the Nucleotide database at NCBI (ncbi.nlm.nih.gov). The Graphics link was used to identify the tRNA genes and their locations, as well as the surrounding non-tRNA genes. The putative tRNA sequences were verified using tRNAscan-SE (Lowe and Eddy, 1997; Lowe and Chan, 2016). Commonalities among the three clades of chloroviruses, as well as their differences were also recorded.

The non-tRNA genes immediately 5’ and 3’ to the tRNA clusters were sought in order to give insight as to whether the 41 chloroviruses acquired their tRNA gene clusters prior to or after their divergence into the three clades. The source of the data was the same NCBI website and it was processed as described above.

### CUB in chloroviruses and their hosts

Because the chloroviruses have GC contents lower than their algal hosts, codon usage was examined for two NC64A viruses and one host, *C. variabilis* NC64A. Geneious 11.1.5 (https://www.geneious.com) was used to determine codon usage for the two viruses and their host.

### Phylogenetic tree construction of three tRNA genes

Three tRNA genes encoding tRNA^tyr^, tRNA^gly^ and tRNA^arg^ were common to all three clades of chloroviruses and were selected for phylogenetic analysis. The tRNA sequences were retrieved from NCBI following BLAST searches. Each set of sequences was aligned by MUSCLE and phylogenies were constructed using maximum parsimony (http://phylogeny.fr). The trees were saved in Newick format and MEGAX was used to produce the final trees (Stecher *et al.,* 2020).

## RESULTS

### Chlorovirus tRNA gene cluster locations

All 41 chloroviruses encode tRNA genes, and the tRNA genes in all 41 viruses were in clusters, usually with intergenic spacers of 1 to ~30 nucleotides. However, in a few cases non-tRNA genes were also present within the tRNA clusters. Collectively, the 14 NC64A viruses had from 7 (virus AR158) to 14 (MA-1E, CvsA1 and CviK1) tRNA genes in their clusters (Table 1A); the 13 SAG viruses had from 7 (NTS-1) to 13 (OR0704.3, Can0619SP, NE-JV-2) tRNA genes in their clusters (Table 1B); and, the 14 Pbi viruses had from 3 (NE-JV-1) to 11 (Fr5L) genes in their tRNA gene clusters (Table1C). If one assumes that the original clusters are the sum of all the tRNAs genes in a clade, the NC64A chlorovirus cluster would consist of 14 tRNA genes; indeed, 3 of the NC64A viruses had all 14 tRNA genes. However, the sum of the SAG viruses was 18 tRNA genes, but the largest number of tRNA genes of any one virus was 13, and the sum of the Pbi viruses was 15 tRNA genes, but the largest number of tRNA genes of any one virus was 11 tRNA genes. Some of the tRNA genes likely represent gene duplications. For example, all 41 chloroviruses had 2 - 4 tRNA^asn^ genes, whose encoded tRNAs recognize the same codon AAC.

**Table 1.**
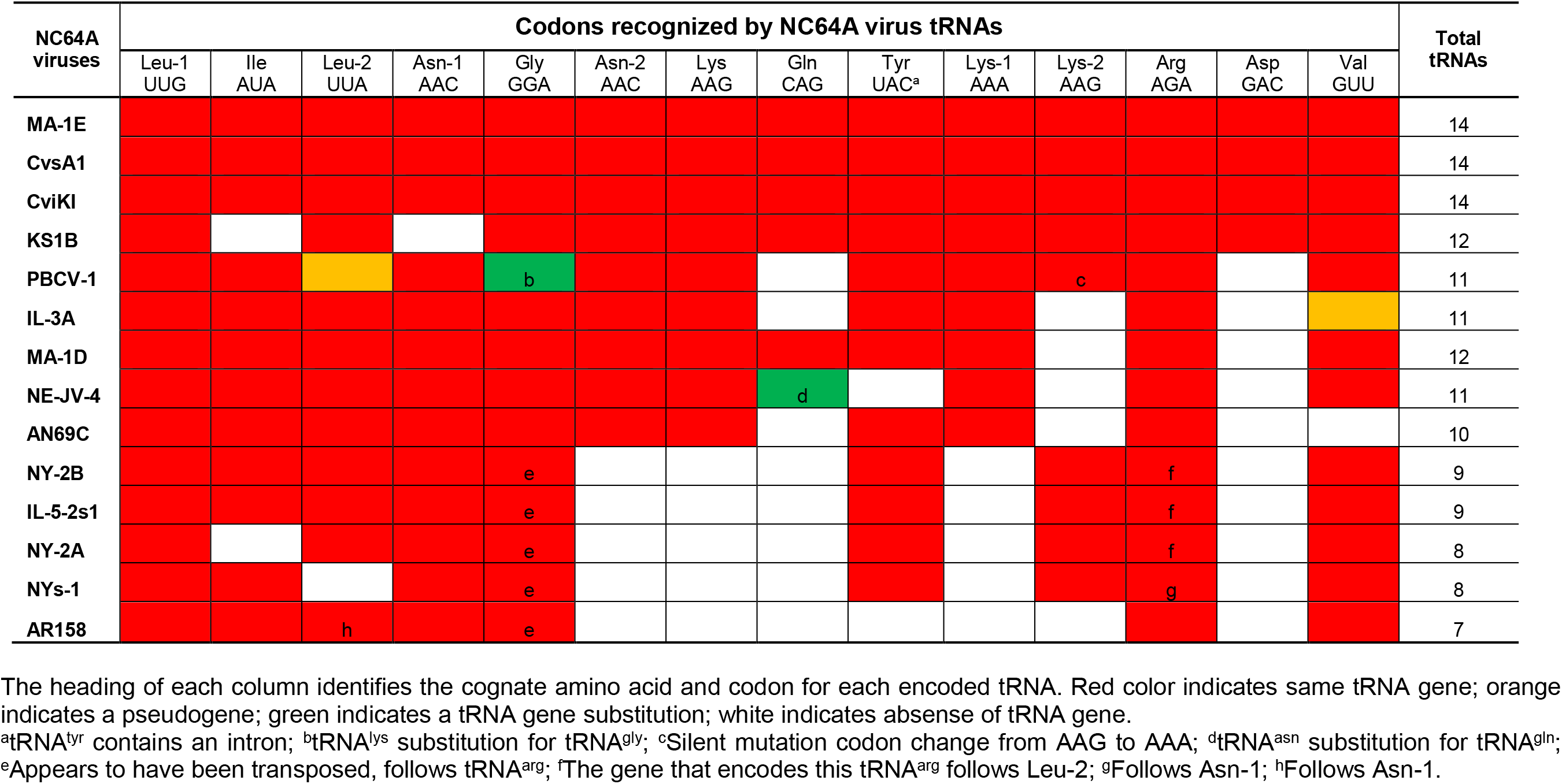
**A)** The order of clustered tRNA genes from NC64A viruses. **B)** The order of clustered tRNA genes from SAG viruses. **C)** The order of clustered tRNA genes from Pbi viruses.

**Table 1B.**
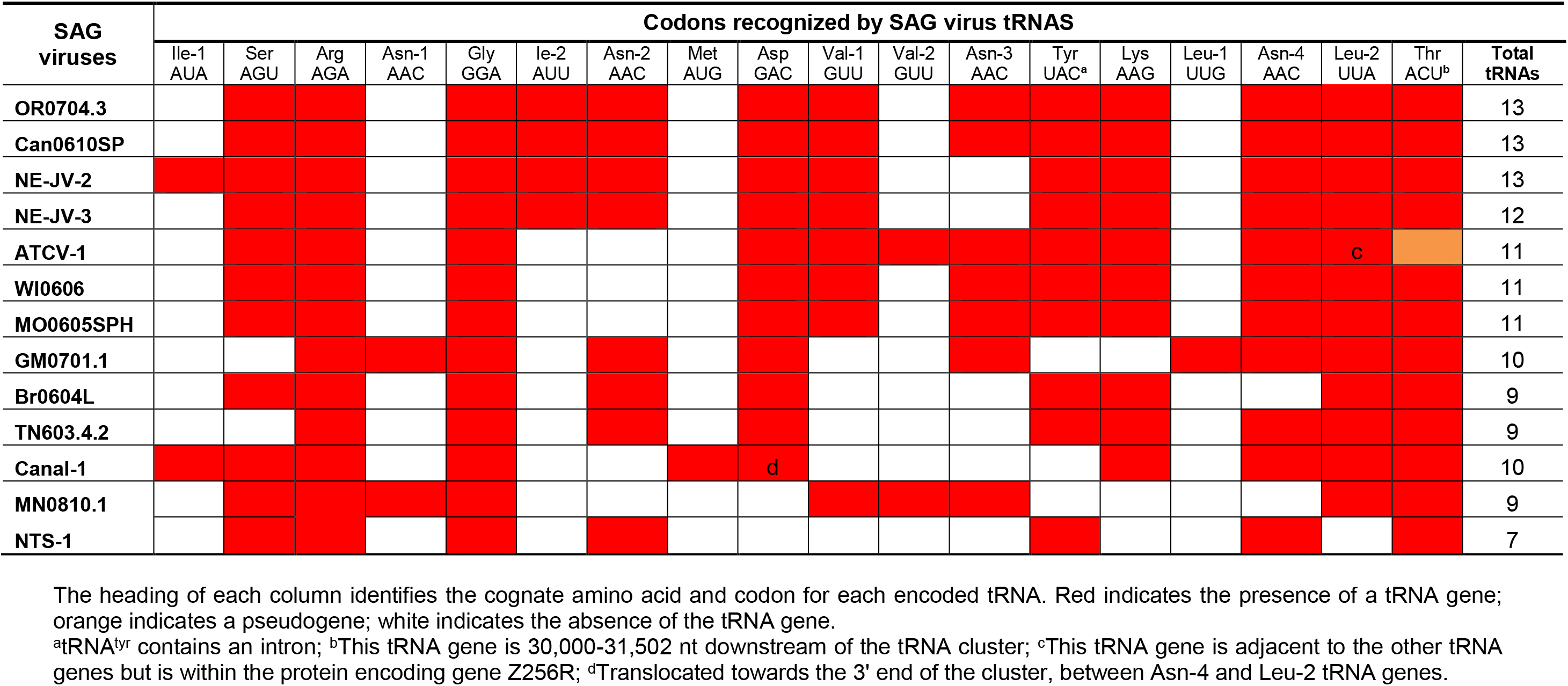
The order of clustered tRNA genes from SAG viruses.

**Table 1C.**
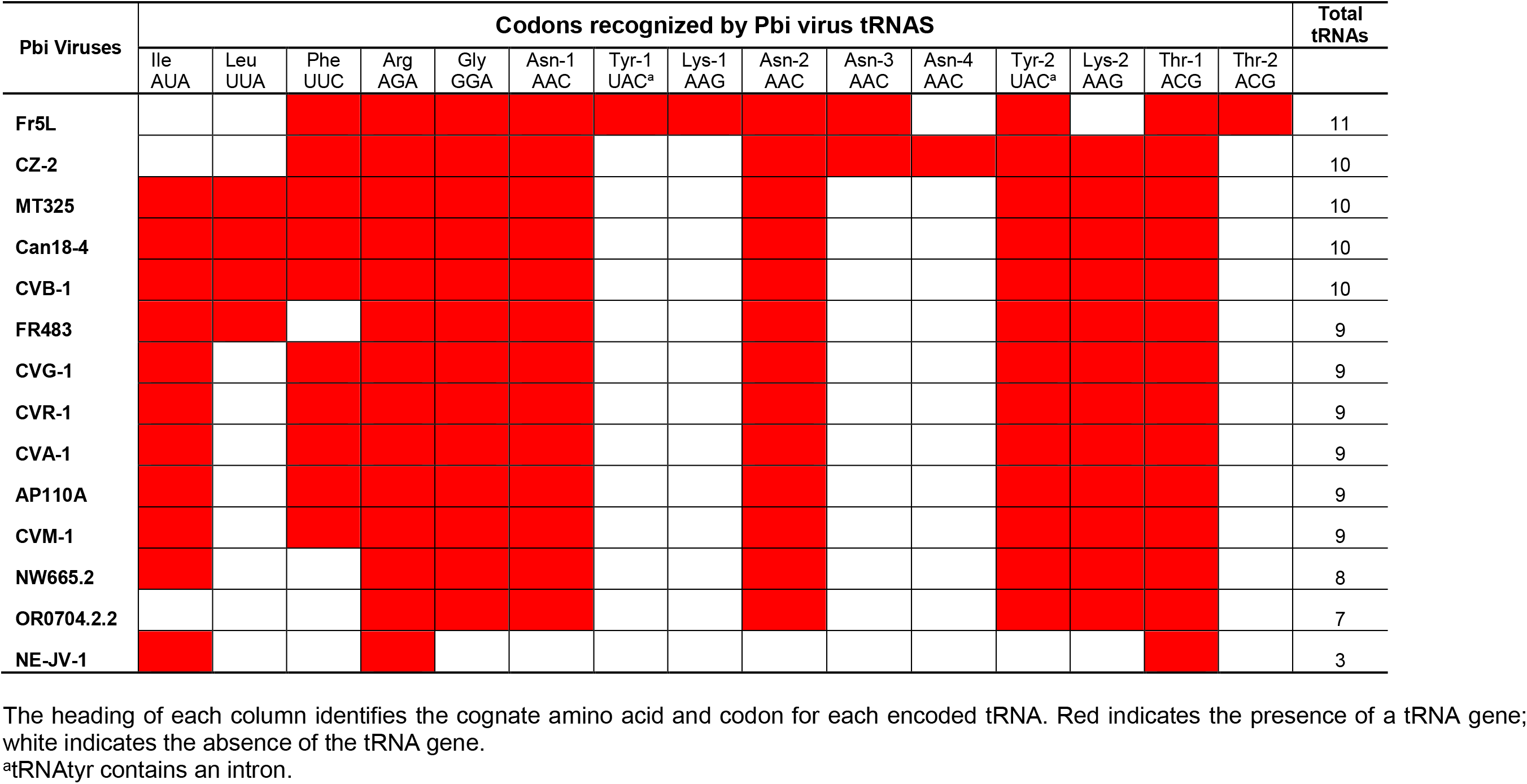
The order of clustered tRNA genes from Pbi viruses.

The order of the chlorovirus tRNA genes within a cluster is also reported in the three heat map tables (Tables 1A-C). In general, there was synteny of tRNA genes among the viruses within a clade, but not between clades. (The exceptions to synteny within a clade are noted in the footnotes of Tables 1A-C.) Seven tRNA genes were common to one or more members of all three chlorovirus clades, although not present in every virus species within any of the three clades (Table 2). Two tRNA genes were unique to the NC64A viruses, 3 tRNA genes were unique to the Pbi viruses and 4 tRNA genes were unique to the SAG viruses. Three additional tRNA genes were common to one or more members of the NC64A and SAG clades. The tRNA^arg^ gene was found in all 41 chloroviruses. Thirty-nine of the 41 chloroviruses had the tRNA^gly^ gene. It is interesting to note that the position in order of encoded tRNAs in the cluster from the 5’ end of the tRNA^gly^ gene was invariant in all 39 viruses.

**Table 2.**
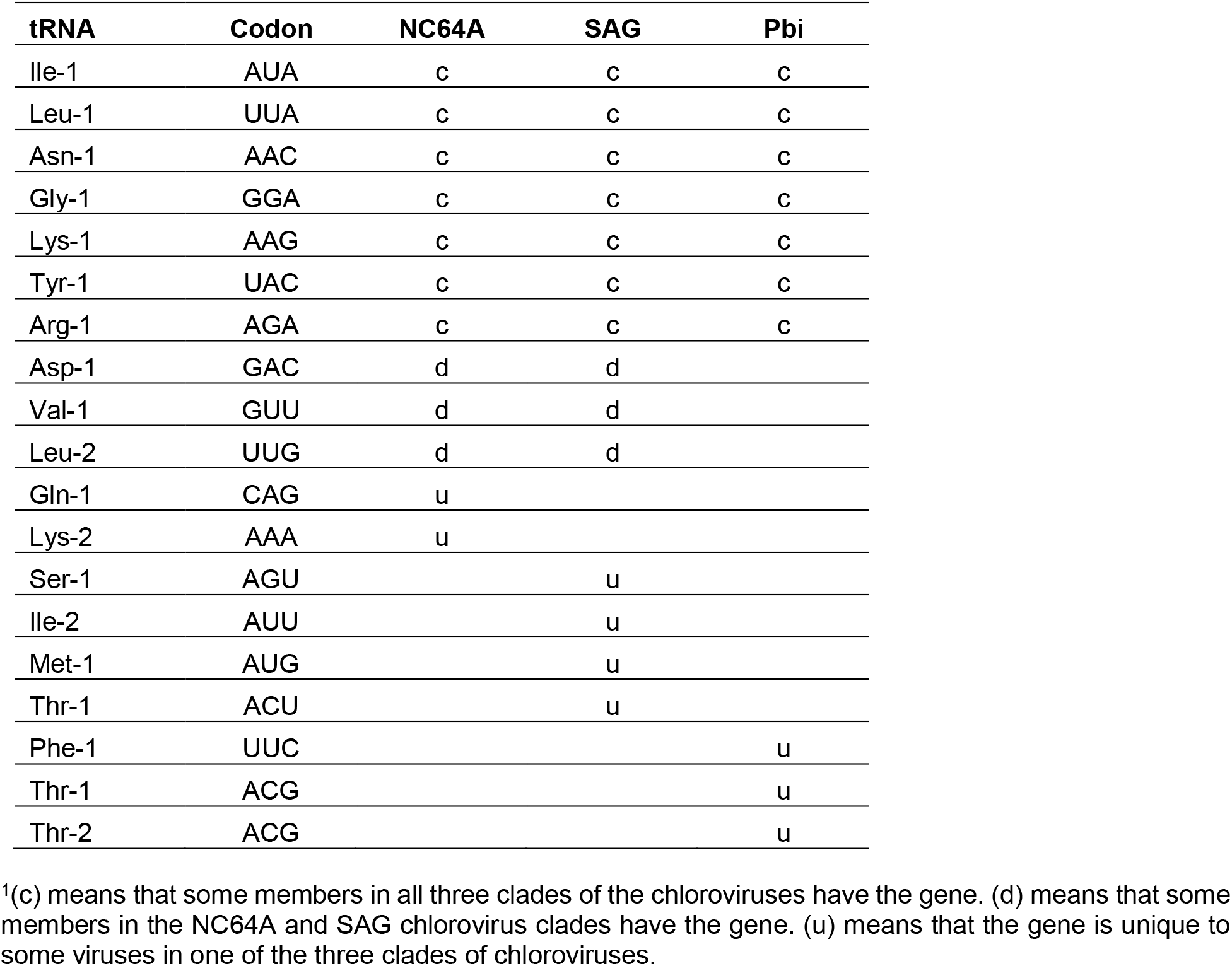
tRNA genes common to one another and unique to each chlorovirus clade^1^.

Differences and similarities in tRNA gene content were observed among individual chloroviruses within each of the three clades of chloroviruses. In some chloroviruses not all of the tRNA genes within a gene cluster were immediately contiguous to one another, leading to interrupted tRNA-gene clusters in which one or several non-tRNA genes were interspersed within the tRNA-gene cluster. This was especially true for the Pbi viruses in which at least four member viruses had interrupted tRNA-gene clusters (Table S2C).

The tRNA clusters in the NC64A and Pbi viruses were located near the center of their genomes except for the Pbi virus NE-JV-1, which was located in the last third of its genome; NE-JV-1 is unusual in many other aspects (Jenniard *et al*., 2013). The tRNA clusters in all of the SAG viruses were located in the first third of their genomes. It is interesting to note that all 13 SAG viruses had a tRNA^thr^ gene located ~30kb beyond the 3’ end of the tRNA gene cluster, placing the tRNA^thr^ genes near the center of their respective genomes.

### Viral tRNAs help to overcome host CUB

Because nuclear genomic codon usage data are available for *C. variabilis* NC64A and its viruses, we were able to compare their respective codon usages (Fig 1 and Table 3; all codons are reported in Fig S1A, S1B). Codon usage was available for 13/14 NC64A viruses, but only two NC64A viruses were included herein as representatives. CUB favoring codons with high GC content were noted in the host alga *C. variabilis* NC64A whose genome is 67% GC (green bars in Fig 1), while the two NC64A viruses, PBCV-1 and AN69C, have GC contents of 40% (Jeanniard et al., 2013) and a CUB favoring AU (Table 3, ratio of codon frequencies comparing virus usage to host usage). For example, in the standard universal code there are four codons for the amino acid alanine, differing only in the third (3’) base of the codon. In *C. variabilis* NC64A the two codons whose third base was C or G (GCC and GCG) were the most common, while the two most common in PBCV-1 and AN69C ended in A and U (GCA and GCU) (Table 3). A parallel example occurs in the usage of the two synonymous codons for glutamic acid; GAG was almost exclusively used by *C. variabilis* NC64A, while GAA was the most common in the two NC64A viruses. The same was true for all amino acids encoded by two synonymous codons (asparagine, aspartic acid, cysteine, glutamic acid, glutamine, histidine, lysine, phenylalanine and tyrosine), as well as other amino acids with more than two synonymous codons. Furthermore, the isoleucine codon AUA (AU-rich) occured in 2.44% of all PBCV-1 codons, while it occured in only 0.20% of codons in *C. variabilis* NC64A, a 12-fold difference (Table 3); hence, in the virus, natural selection appears to have favored the retention of the gene that encodes the cognate tRNA for this codon. Likewise, the lysine codon AAA (AU rich) occured in 4.7% of all PBCV-1 codons, but in only 0.25% of host codons, nearly a 20-fold difference. The leucine codon UUA was the rarest codon used by *C. variabilis* NC64A (0.06% of all codons) while it was a moderately common codon used by PBCV-1 (1.39% of all codons) (Table 3). As a final example, there are six synonymous codons for the amino acid arginine, but only one codon that had one guanine or cytosine (AGA), while the other five codons had a minimum of two guanines and/or cytosines. 1.43% of all viral codons were AGA for arginine, but only 0.25% of *C. variabilis* NC64A codons were AGA. Indeed, the two arginine codons with three guanines and cytosines (CGC and CGG) were the two most common in *C. variabilis* NC64A (3.22% and 2.11% respectively), while the same two codons in the two viruses averaged 0.75% and 0.70%, respectively.

**Table 3.**
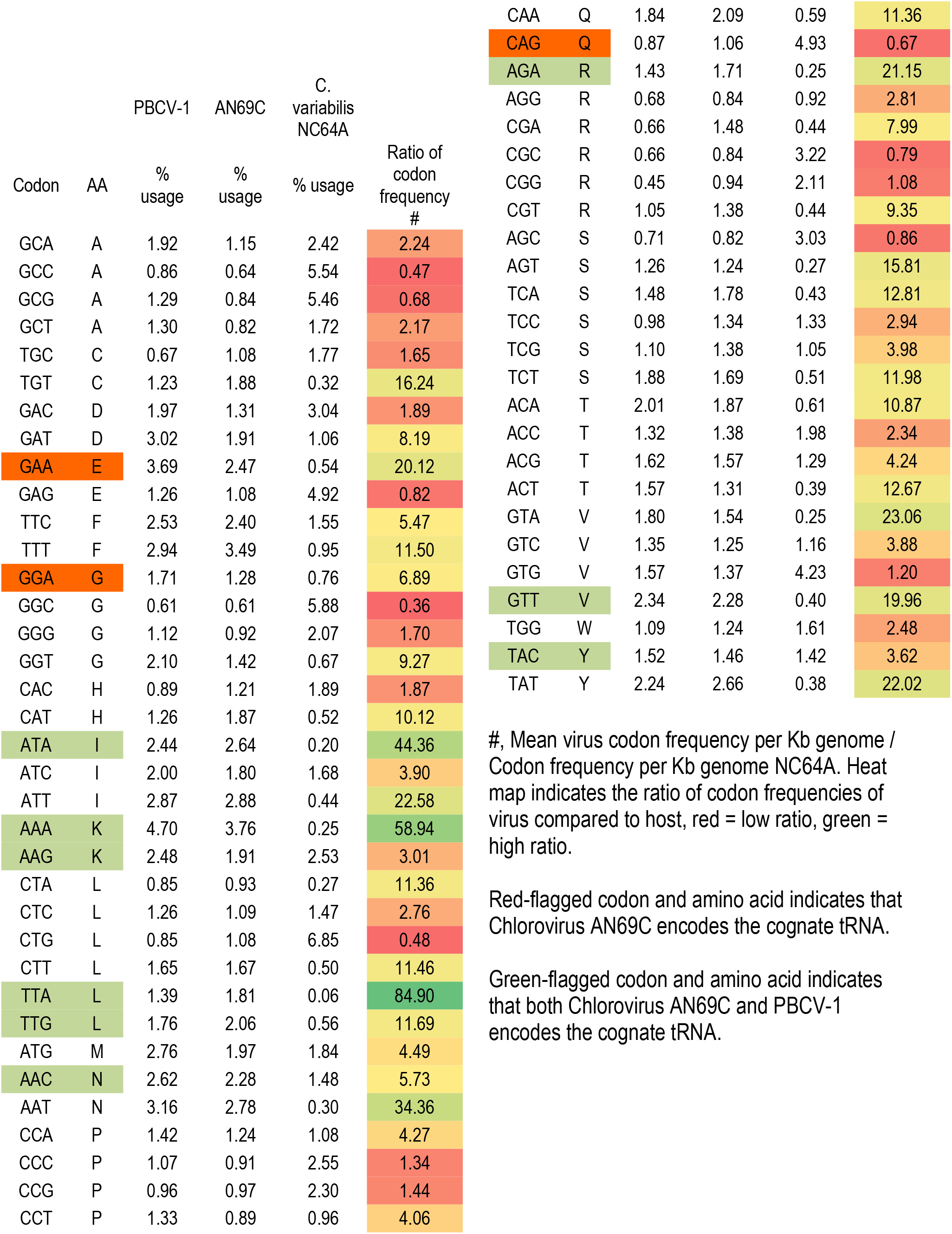
Codon frequency use comparison: virus to host

**Figure 1.**
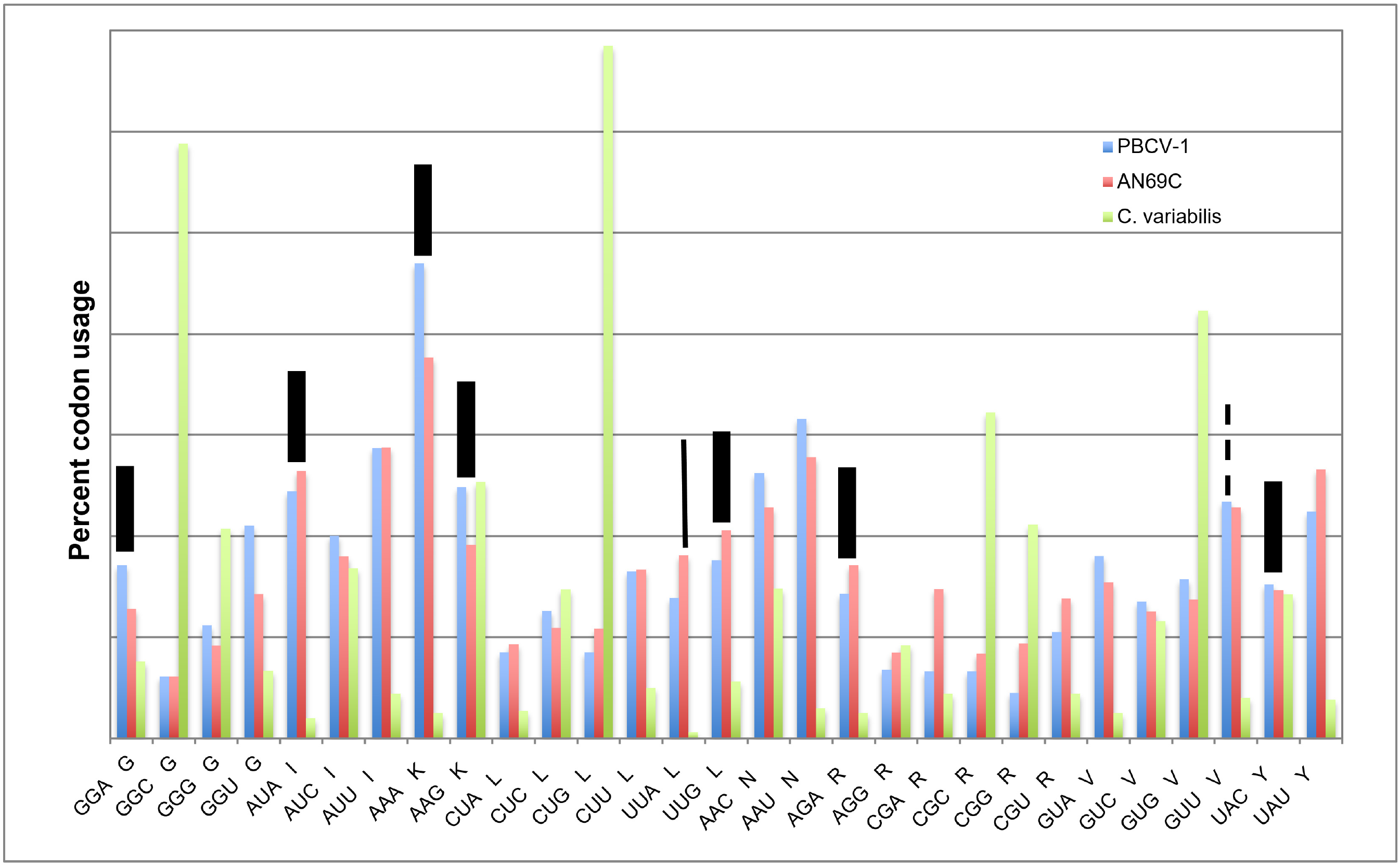
Comparison of the frequency of codon usage recognized by tRNAs encoded by PBCV-1 and AN69C, compared with their host *C. variabilis* NC64A. Wide bars denote the codons recognized by tRNAs encoded by PBCV-1 and AN69A. Narrow solid bar denotes codon recognized by tRNA encoded by AN69C but not PBCV-1. Narrow dash bar denotes codon recognized by tRNA encoded by PBCV-1 but not AN69C.

### Acquisition of chlorovirus tRNA clusters

One question we wished to address was: did the three chlorovirus clades independently acquire their tRNA clusters by horizontal gene transfer (HGT) or was the tRNA gene cluster acquired by the MRCA, i.e., the last common ancestor prior to splitting into the three clades? We used several lines of inquiry to address this question. First, as described above, the three clades of chloroviruses had seven tRNA genes in common with one another (Table 2), including 2 - 4 tRNA^asn^ genes, whose encoded tRNAs recognize the codon AAC. There were, however, considerable differences in composition and order between clades, unlike the similar composition and synteny within each of the three clades (Tables 1A-C).

The second line of inquiry focused on the protein-encoding genes that border the 5’ and 3’ sides of the tRNA clusters. Supplementary Tables S2A-C report the commonness of protein-encoding genes (orthologous genes) within each of the three clades, but the protein-encoding genes surrounding the tRNA clusters of each clade were not orthologous across the three clades.

A third line of inquiry was to examine phylogenetic tree constructs of three tRNA genes common to all three clades of chloroviruses: tRNA^gly^, tRNA^arg^ and tRNA^tyr^ (Fig 2A, 2B and 2C, respectively). Most of the chloroviruses tended to cluster within their clade, but there were a number of exceptions, including the presence of subclades, as will be discussed below.

**Figure 2A.**
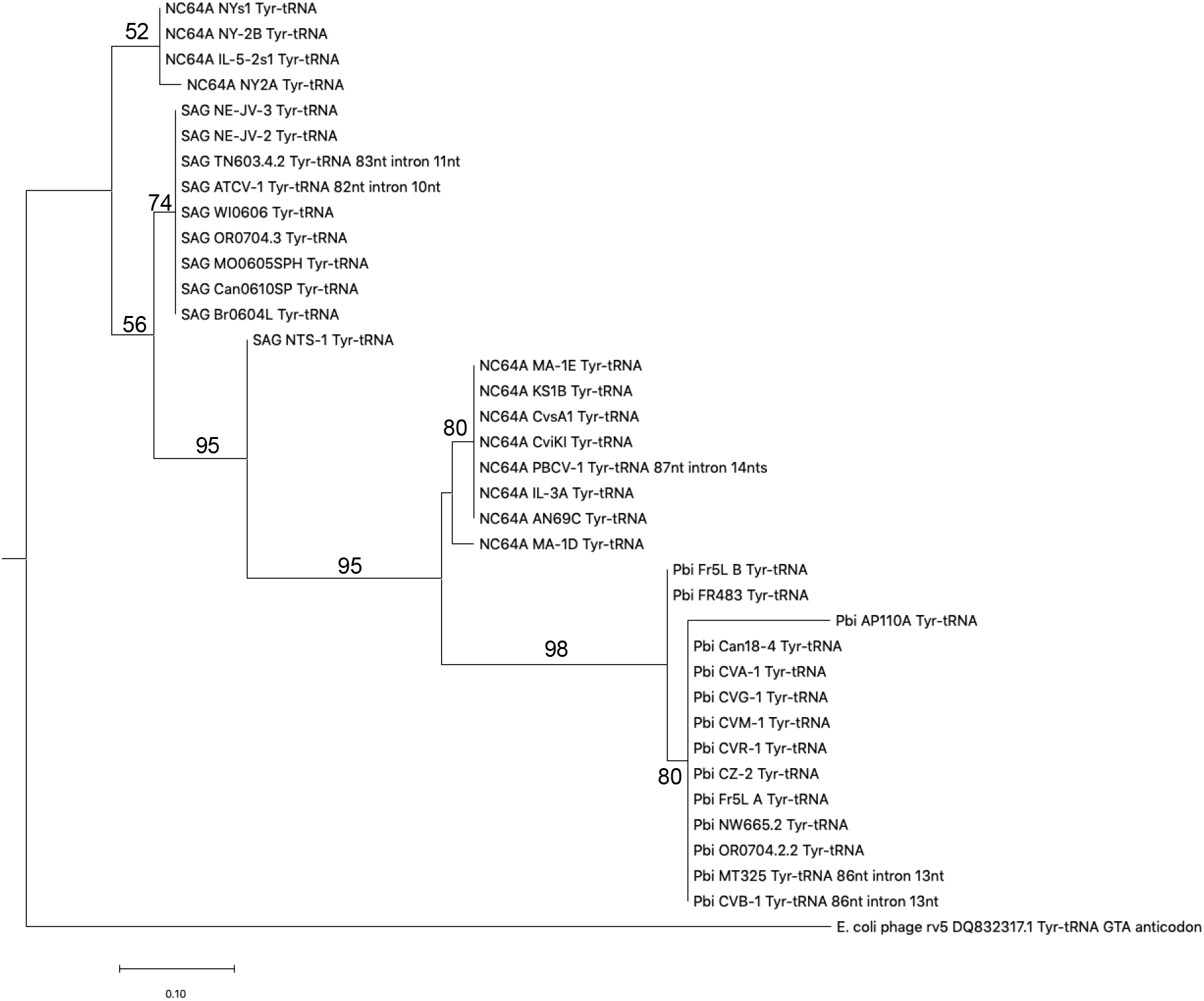
**A**) Phylogenetic tree of tRNA^tyr^ genes from 35 chloroviruses representing all three clades, NC64A, Pbi and SAG. Six chloroviruses lacked the tRNA^tyr^ gene. The tRNA^tyr^ gene from *E. coli* phage rv5 was used as the outgroup. One subclade of NC64A viruses are more similar to SAG viruses, while the other subclade of NC64A viruses are more similar to the Pbi viruses. **B**) Phylogenetic tree of tRNA^gly^ genes from 39 chloroviruses representing all three clades, NC64A, Pbi and SAG. Two chloroviruses lacked the tRNA^gly^ gene. The tRNA^gly^ gene from cyanophage NATL1A was used as the outgroup. One NC64A subclade is more similar to SAG viruses than the other subclade of NC64A viruses. Two SAG viruses and two Pbi viruses are more similar to a second subclade of NC64A than they are to viruses in their own clades. **C**) Phylogenetic tree of tRNA^arg^ genes from all 41 chloroviruses representing all three clades, NC64A, Pbi and SAG. The tRNA^arg^ gene from enterobacteria phage EU330206.1 was used as the outgroup. One NC64A subclade and two SAG viruses are more similar to Pbi viruses than they are to viruses in their own clade. Bootstrap values greater than 50 are reported. The sequences were aligned with MUSCLE and the trees were constructed using the maximum likelihood algorithm.

**Figure 2B.**
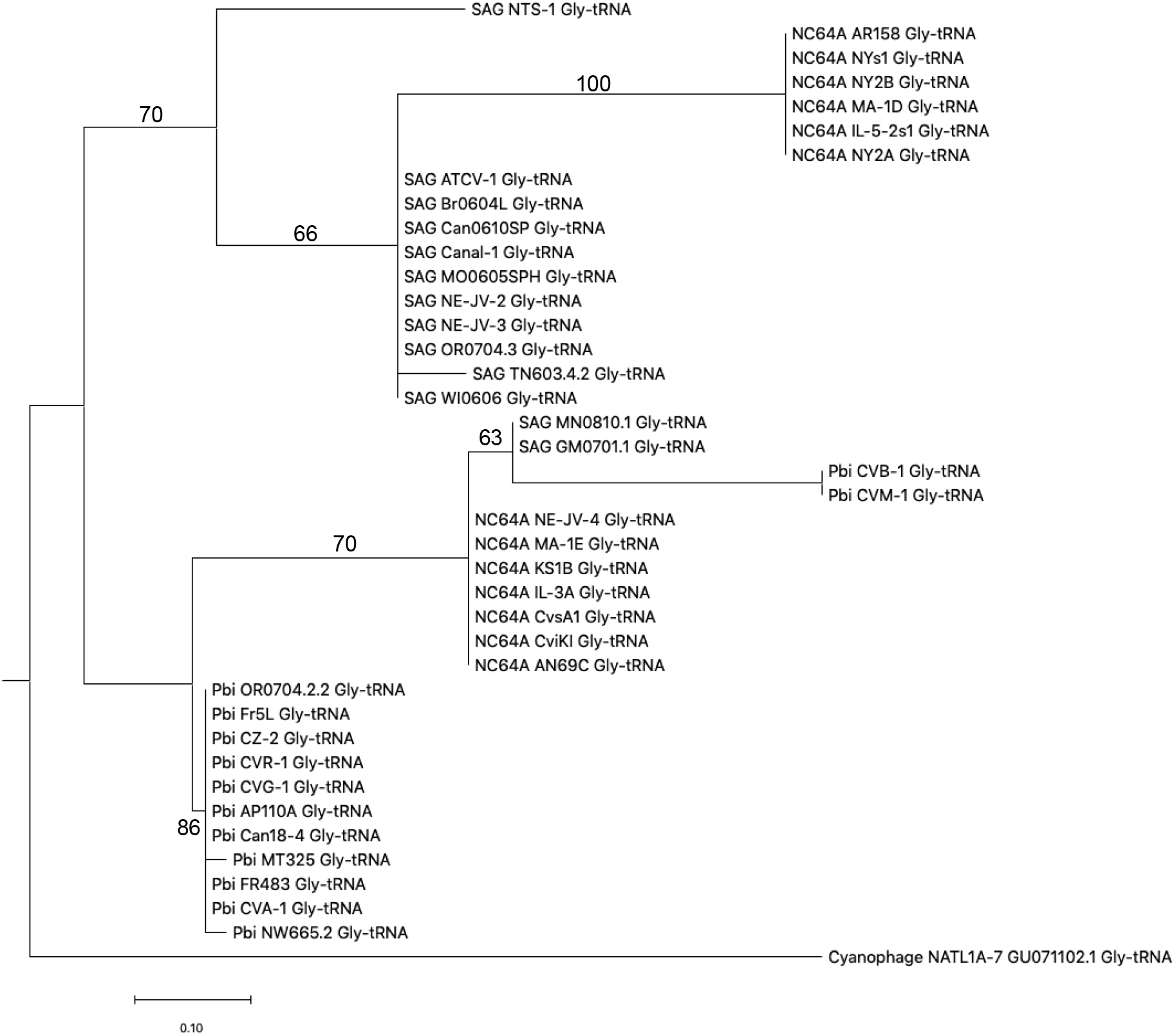
Phylogenetic tree of tRNA^gly^ genes from 39 chloroviruses representing all three clades, NC64A, Pbi and SAG. Two chloroviruses lacked the tRNA^gly^ gene. The tRNA^gly^ gene from cyanophage NATL1A was used as the outgroup. One NC64A subclade is more similar to SAG viruses than the other subclade of NC64A viruses. Two SAG viruses and two Pbi viruses are more similar to a second subclade of NC64A than they are to viruses in their own clades. Bootstrap values greater than 50 are reported. The sequences were aligned with MUSCLE and the trees were constructed using the maximum likelihood algorithm.

**Figure 2C.**
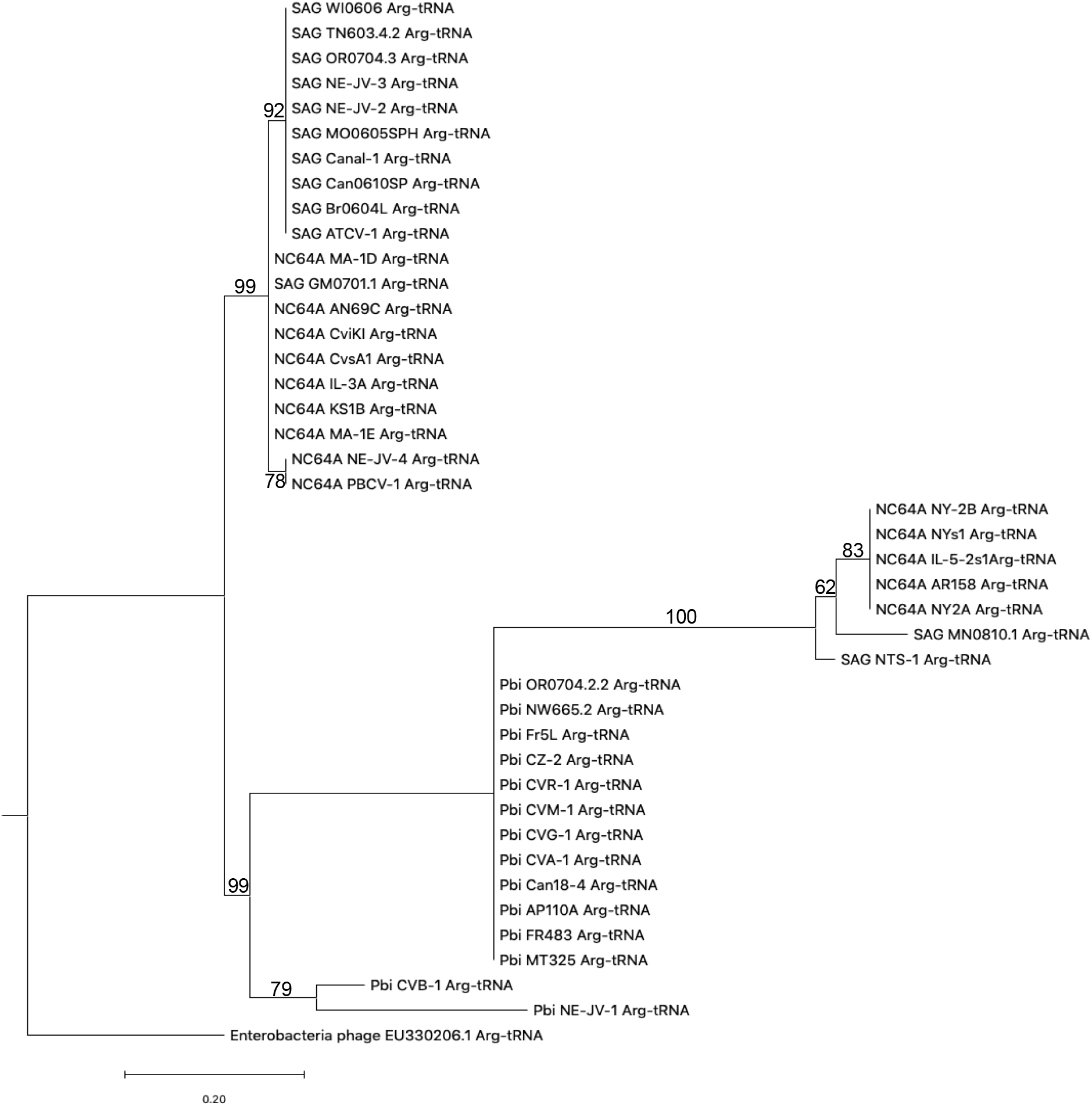
Phylogenetic tree of tRNA^arg^ genes from all 41 chloroviruses representing all three clades, NC64A, Pbi and SAG. The tRNA^arg^ gene from enterobacteria phage EU330206.1 was used as the outgroup. One NC64A subclade and two SAG viruses are more similar to Pbi viruses than they are to viruses in their own clade. Bootstrap values greater than 50 are reported. The sequences were aligned with MUSCLE and the trees were constructed using the maximum likelihood algorithm.

A fourth line of inquiry focused on the tRNA^tyr^ gene found in 34/41 chloroviruses representing all three clades, all 34 of which had an intron (Table S3). We examined the locations and sequences of the introns and found that the introns were located in identical positions in all the tRNA^tyr^ genes, one nucleotide from the anticodon (3’ direction). While the intron lengths and sequences were nearly identical within a clade, there were slight differences in length between clades: NC64A 13-14 nt; Pbi 13 nt; SAG 10-11 nt in the three clades.

## DISCUSSION

### Structural features of the chlorovirus tRNA clusters

The results presented in this paper indicate that all 41 chloroviruses, representing three clades, had clusters of tRNAs with intergenic spacers usually of 1 to ~30 nucleotides. Collectively, the chloroviruses encoded a total of 410 tRNAs of which there were 17 different tRNAs for 14 different amino acids (3 synonymous codons). The large numbers of viral tRNA genes in the chloroviruses should not be surprising because Morgado and Vicente (2019) report a positive correlation between viral genome length and the number of tRNA genes in viruses. In addition, they found that tRNA gene clusters are more common in viruses with larger genomes than in those with smaller genomes. All the individual chlorovirus-encoded tRNAs lacked the 3’ terminal three nucleotides (CCA) necessary in order to be aminoacylated. Since fully functional tRNAs were reported for the NC64A chlorovirus CVK2 (Nishida *et al*., 1999), the chlorovirus tRNAs, like tRNAs from cellular organisms (Rak *et al*., 2018), must either use an unidentified virus enzyme(s) or the host tRNA nucleotidytransferase to add the CCA nucleotides prior to tRNA aminoacylation for functionality.

Due to the short intergenic spacers between clustered chlorovirus tRNA genes, we suspect that the tRNA gene clusters are transcribed as one transcript. Indeed, Nishida *et al*., (1999) reported that chlorovirus CVK2 transcribes its tRNA gene cluster of 14 tRNAs into one transcript; furthermore, they reported that the RNA transcript was precisely processed into individual tRNA species by either some unknown virus-encoded or host-encoded RNase. In this regards, PBCV-1 encodes a functional RNase III enzyme (Zhang *et al*., 2003) but its role in virus replication is unknown.

Other viruses in the *Phycodnaviridae* family also have tRNA gene clusters. *Micromonas pusilla* virus 12T encodes six tRNA genes, five of which are clustered (NC_020864.1). The sixth tRNA gene, tRNA^thr^, is an orphan ~30kb beyond the tRNA^leu^ gene, the 3’ member of the tRNA cluster. The position of these two tRNA genes and the distance between them is the same as in the chlorovirus SAG clade; i.e., there was a 30kb gap between the genes that encode tRNA^leu^ and tRNA^thr^. The virus 12T also encodes a tRNA^tyr^, as do the chloroviruses, but unlike the chloroviruses, this *Micromonas* virus does not have an intron in its tRNA^tyr^ gene. On the other hand, *Ostreococcus lucimarinus* virus 7 (OlV7) (Derelle *et al*., 2015; KP874737), which has five tRNA-clustered genes, encodes a tRNA^tyr^ that does have a 15 nt intron located in the same position as the chloroviruses. In fact, the tRNA genes in the *Micromonas* and *Ostreococcus* virus clusters are similar to some of the tRNA gene clusters in the NC64A viruses, suggesting the two groups of viruses might have a common evolutionary ancestor. Thus, while not all viruses of the *Phycodnaviridae* family were examined, tRNA clusters appear to be common in this family.

### Functional features of the chlorovirus tRNA clusters

The chlorovirus-encoded tRNAs were evaluated to determine if they help the viruses overcome the CUB ascribed to the host (Fig 1 and Table 3). That is, most of the chlorovirus encoded tRNAs had codons favoring AU, whereas, the host codons had a GC bias. Nishida *et al*. (1999) reached a similar conclusion for the chloroviruses. Therefore, we conclude that the chloroviruses partially solved the CUB problem by encoding some tRNAs that support a virus AU bias. At least some of the differences in tRNA gene content between clades is likely due to the evolutionary pressures of utilizing three different hosts. We also observed unexplained differences within clades. Only tRNA^arg^ was common to all 41 viruses. In the SAG viruses the tRNA^thr^ gene that recognized ACU was located ~30kb downstream of the clusters. The Pbi virus NE-JV-1 is very odd in many aspects (Jeanniard *et al*., 2013), including that it encoded only three tRNAs and the location of its tRNA cluster was far downstream of all the 40 other viruses. So, with the exclusion of NE-JV-1, 40/40 chloroviruses had genes that encoded tRNA^asn^ and tRNA^gly^, while 37/40 encoded tRNA^lys^.

However, not all tRNA members of a cluster assist in overcoming the CUB of the host. For example, PBCV-1 encoded 11 tRNAs but only seven help it to overcome CUB; likewise, AN69C encoded ten tRNAs but only eight favor its own CUB (Table 1A). Thus, we speculate that the cluster as a unit is under natural selection because some of the tRNA genes in the clusters are neutral and do not bestow any positive selective advantage. A reasonable explanation for these observations is that the neutral tRNA genes are preserved by natural selection as hitchhikers due to their close linkage to tRNA genes that help the viruses overcome the CUB of the host. To illustrate this point, all three clades of chloroviruses had 2-4 tRNA^asn^ genes whose tRNAs only recognize the codon AAC; none of the 41 chloroviruses encoded a tRNA^asn^ that would recognize the alternate codon AAU, which was 10X more common in the two viruses than in *C. variabilis* (Table 3), and which would presumably enhance viral protein translation. Of no surprise, the most common of the two Asn codons in *C. variabilis* is AAC. The presence of the AAC codon cognate tRNA^asn^ genes, which appear to be neutral in benefit to all 41 chloroviruses, suggests that they were acquired in the MRCA tRNA cluster due to an evolutionary accident. That is, their presence appears to be an artifact of evolutionary history by which some neutral tRNA genes were acquired in the tRNA cluster along with selectively advantageous genes by some mechanism, such as HGT. (This kind of event is similar to the frozen accident concept first proposed by Crick (1968).) In the case at hand, neutral tRNA^asn^ genes may be maintained in the tRNA gene clusters seemingly due to their tight linkage to selectively advantageous viral tRNA genes in the cluster. The same argument might explain the presence in PBCV-1 of the tRNA^lys^ gene for the cognate codon AAG. PBCV-1 encoded both of the tRNAs genes that recognize the two lysine codons, AAG and AAA. However, the AAA codon was >20X more common in PBCV-1 than in *C. variabilis*, while AAG was the most common lysine codon in *C. variabilis*; hence, the former appears to be maintained by positive selection while the latter appears to provide a neutral benefit to PBCV-1.

Thirty-nine of the 41 chloroviruses encode another putative enzyme involved in codon usage, tRNA isoleucine lysidine synthase (TilS). The methionine codon AUG is normally recognized by its cognate tRNA^met^ with the 3’ UAC 5’ anticodon. The TilS enzyme ligates lysine to the cytidine in the 5’ position of the tRNA anticodon; this modified cytidine becomes lysidine, which is complementary to adenine in the 3’ position of the codon, rather than guanine (Fig 3). As such, this modified tRNA then behaves as a tRNA^ile^ and recognizes the isoleucine AUA codon (Nakanishi et al., 2005; Suzuki and Miyauchi, 2010). The AUA codon was 12-13 times more common in the two NC64A viruses than in the host, *C. variabilis* (Table 3). Thus, we suspect this enzyme provides one additional mechanism that helps the viruses overcome CUB of the host by enabling more efficient viral protein synthesis, diminishing the chances of a ribosome stall during elongation when AUA codons are encountered. In addition, all chloroviruses encode a homolog of translational elongation factor 3 (EF-3) (Jeanniard et al., 2013). EF-3 plays a role in optimizing the accuracy of mRNA decoding at the ribosomal acceptor site during protein synthesis in fungi (Belfield and Tuite, 1993; Belfield *et al*., 1995); EF-3 has been reported recently in algae (Mateyak *et al*., 2018). The role this putative enzyme plays in chlorovirus translation is unknown.

**Figure 3.**
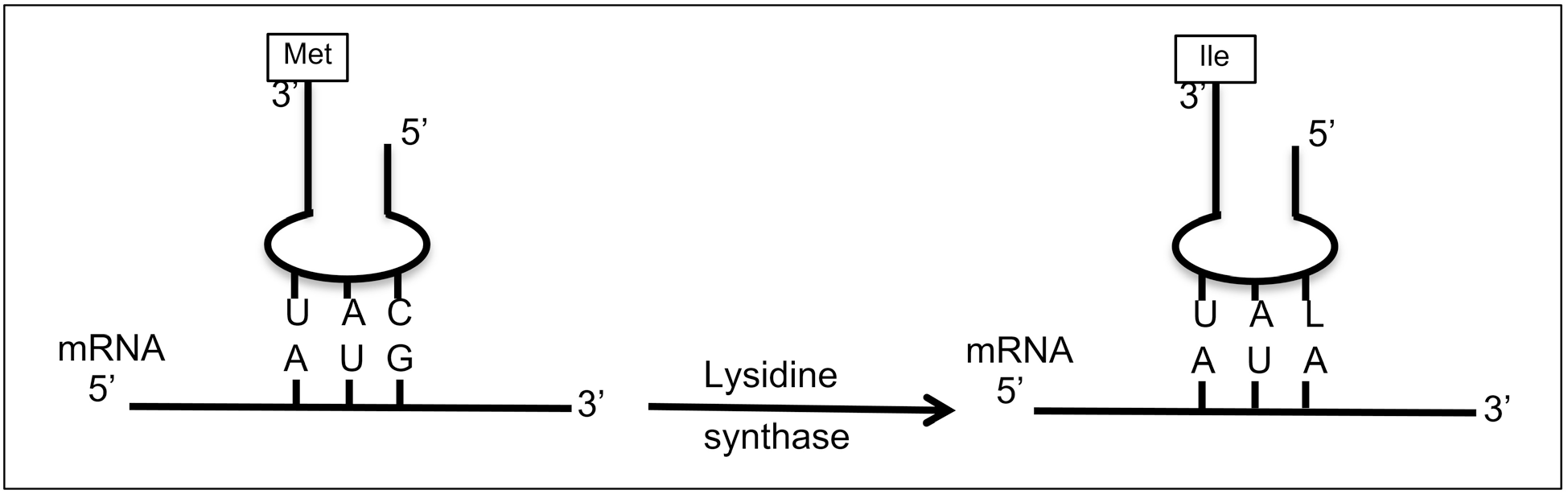
Simplified diagram of tRNA anticodon base pairing with the mRNA codon. The tRNA^met^ anticodon 3’ UAC 5’ recognizes the codon 5’ AUG 3’ (left figure). The enzyme tRNA ile lysidine synthase (TilS) attaches lysine to the 5’ cytosine of the tRNA, which becomes lysidine. Lysidine base pairs with adenine. As such, the modified tRNA now recognizes the isoleucine codon AUA.

### Evolutionary history of the chlorovirus tRNA clusters

The origin(s) of the chlorovirus tRNA clusters is intriguing, i.e., was there a single HGT event in the MRCA prior to the evolution of the chlorovirus clades, or did individual HGT events occur independently in each clade post-MRCA? A previous study suggests that the chloroviruses had a common host prior to their current three hosts (Jenniard *et al*., 2013). Support for the argument that the viruses may have acquired the tRNA clusters shortly after they separated into the three clades, includes: i) while the three clades of chloroviruses had some tRNA genes in common with one another, there were differences in the composition and gene order among the clades, unlike the somewhat uniform composition and order within each independent clade. ii) The protein-encoding genes that border the 5’ and 3’ sides of the tRNA clusters were orthologous for the viruses within each of the three clades but differed across the three clades.

In contrast, four lines of evidence favor the notion that the viruses acquired the tRNA cluster by a single HGT event in the MRCA prior to the evolution of all three clades of chloroviruses. i) 34/41 chloroviruses had a tRNA^tyr^ gene and in all 34 cases the gene had a tRNA intron (Schmidt and Matera, 2019), which was located in an identical position among the 34 viruses representing all three clades, suggesting that the intron was inserted as a single event. While the lengths and sequences of the introns were nearly identical within a clade there were small differences in length between clades: NC64A 13-14 nt; Pbi 13 nt; SAG 10-11 nt (Table S3). Point mutations and indels over evolutionary time could explain the differences within and between the clades. ii) All three clades had several tRNA genes in common, as described above and seen in Tables 1A-C. Perhaps the most interesting is the tRNA^asn^ gene, which occured in 2-4 copies in 40/40 chloroviruses (excluding NE-JV-1). Every single tRNA^asn^ recognizes the codon AAC, but none of the 40 viruses encoded a tRNA^asn^ cognate for the codon AAU, which was the most common codon in the viruses and least common in the host (Table 3). Independent acquisition of the same neutral tRNA gene by each of the three clades seems unlikely. iii) The third line of support can be seen in Figures 2A, 2B and 2C, which display the phylogenetic tree constructs of tRNA^gly^, tRNA^tyr^ and tRNA^arg^ genes, genes that were common to all three clades. While most of the viruses assorted within their respective clades, it was the exceptions that support a single HGT event hypothesis. For example, the tRNA^tyr^ genes of four of the NC64A viruses were more similar to the SAG viruses than to other NC64A viruses (Fig 2A). Likewise, for the tRNA^gly^ gene, half of the NC64A viruses were more similar to SAG viruses, while the other half of the NC64A viruses were more similar to Pbi viruses (Fig 2B); furthermore, there were two Pbi and two SAG viruses that were more similar to a subclade of NC64A viruses than to other members of their own clade. iv) The fourth line of support involves the tRNA^thr^ gene that was found in all of the SAG and Pbi viruses. In all 14 of the Pbi viruses, this gene was the most 3’ member of the cluster, but for all 13 SAG viruses, the tRNA^thr^ gene was located ~30kb beyond the 3’ end of the tRNA cluster. One possible explanation is that the tRNA cluster in the SAG viruses was translocated in the 5’ direction, but without the tRNA^thr^ gene, prior to speciation within the SAG clade. Indeed, unlike the NC64A and Pbi viruses whose tRNA clusters were located near the center of their genomes, the tRNA clusters in the SAG viruses were located more towards the 5’ direction of their genomes – i.e., in the first third of their genomes. If a translocation did take place, then the original location of the cluster in the ancestral SAG virus would have been more towards the center of its genome. It is perhaps of note that the average genome size of the NC64A, Pbi and SAG clades is as follows: 326, 321 and 307 kb, respectively (Jenniard *et al*., 2013). A translocation event could not only explain the relocation of the SAG tRNA gene cluster, but also the reduction in genome size of the ancestral SAG virus.

Therefore, we feel that the evidence most strongly supports a single origin for the chloroviruses tRNA gene clusters. Regardless of the origin, it is clear that many evolutionary changes have occurred. The ability of tRNA genes to proliferate is thought to be similar to the mechanism by which mobile elements can lead to intragenomic gene duplications (Velandia-Huerto *et al*., 2016). Duplications and losses are clearly evident among and within all three chlorovirus clades; for example, the Pbi virus CZ-2 had four tRNA^asn^ genes while almost all of the other Pbi viruses had just two copies. As well, the NC64A virus MA-1E had three tRNA^lys^ genes, while some others in the same clade had just one, and AR158 had none. There are other similar examples among the three chlorovirus clades.

Pope *et al*. (2014) implicates a homing endonuclease (HNH) in the generation of tRNA genes in mycobacteriophages. While not immediately adjacent to the tRNA gene clusters, all of the chloroviruses encode at least two putative HNHs (e.g., orthologs of A087R and A422R in PBCV-1). A second source of HNH or other endonucleases might have been from co-infecting viruses that had such genes in their repertoire that generated tRNA gene duplications, losses and translocations within the chloroviruses. Previously, we proposed three potential sources of HGT for the chlorovirus protein-encoding genes that might also explain the tRNA clusters in the chloroviruses: (i) viral host(s), although there are only a few NC64A chlorovirus genes that have likely been acquired from *C. variabilis* NC64A but the viruses probably had at least one other host through evolutionary time; (ii) bacteria, because some of the chlorovirus genes appear to be of bacterial origin; and, (iii) from other host-competing viral species (Jeanniard *et al.*, 2013). Plastids and mitochondria might also have contributed to the viruses via HGT, at least for the tRNA gene clusters, because those organelles have tRNA gene clusters, as well. These results are consistent with the analyses and conclusions of Fan *et al*. (2017) who sequenced the mitochondria and plastids of the three chlorovirus hosts, *C. variabilis*, *C. heliozoae* and *M. conductrix.* Indeed, Margado and Vicente (2019) propose that viruses with tRNA clusters might be the source of dissemination of such clustered structures in the three domains of life. Acquisition of clustered genes appears to be a common occurrence among the chloroviruses; six of the chloroviruses examined by Filée et al. (2008) appear to have acquired genes from both bacteria and eukarotes, in many cases they were in clusters (see Supplementary Material; Additional File 1 in the Filée reference). The chlorovirus genomes appear to have been randomly inserted with those acquired genes. Two of the six chloroviruses that were examined had numerous insertion elements, which could explain the movement of genes between genomes and within genomes. Additionally, gene gangs are conserved clusters of colinear monocistronic chlorovirus genes, some of which have an apparent common origin (Seitzer et al., 2018).

In the larger picture about viruses encoding constituents of the protein synthesis machinery, it is clear that many of the recently discovered giant viruses encode components of the protein synthetic machinery (e.g., Brandes and Linial, 2019). The first giant virus to be discovered, *Acanthamoeba polyphaga mimivirus* (AMPV), encodes four putative aminoacyl tRNA synthetases (aaRS) and six tRNAs (Raoult *et al*., 2004), which, unlike the chloroviruses, are not clustered. One of the four mimivirus aaRSs, the tyrosine tRNA synthetase, has been crystalized and shown to function (Abergel *et al*., 2005). More amazing, two recently described giant Tupanviruses, a close relative of the mimiviruses, encode up to 70 tRNA genes, many in clusters of ~15 tRNAs, and 20 aaRS genes, plus many more protein synthesis genes (Abrahao *et al*., 2018). Thus, these newly discovered giant viruses are apparently solving the CUB issues by encoding several elements of the protein synthetic machinery. It will be exciting to find out if all of these putative proteins have their predicted activities.

In summary, the two hosts for the chloroviruses that have been genomically sequenced have a high GC content (67%) and natural selection tends to favor GC-rich codons, whereas in the chloroviruses natural selection favored AU-rich codons. The viruses appear to overcome the CUB of their hosts by maintaining a cluster of tRNA-encoding genes that favor cognate codons richer in AU. However, the evolutionary events have differed among the viruses, even within the same clade, because very few of the tRNAs were conserved among all of the chloroviruses. We are aware that tRNAs are turning out to play other roles in cells (e.g., Lyons et al., 2018) and so it is always possible that the viral encoded tRNAs have some other functions.

## ACKNOWLEDGEMENTS

This research was supported by the National Science Foundation under Grant No. 1736030 and the University of Nebraska-Lincoln Agricultural Research Division and the Office of Research and Economic Development.

---

**Table S1.**
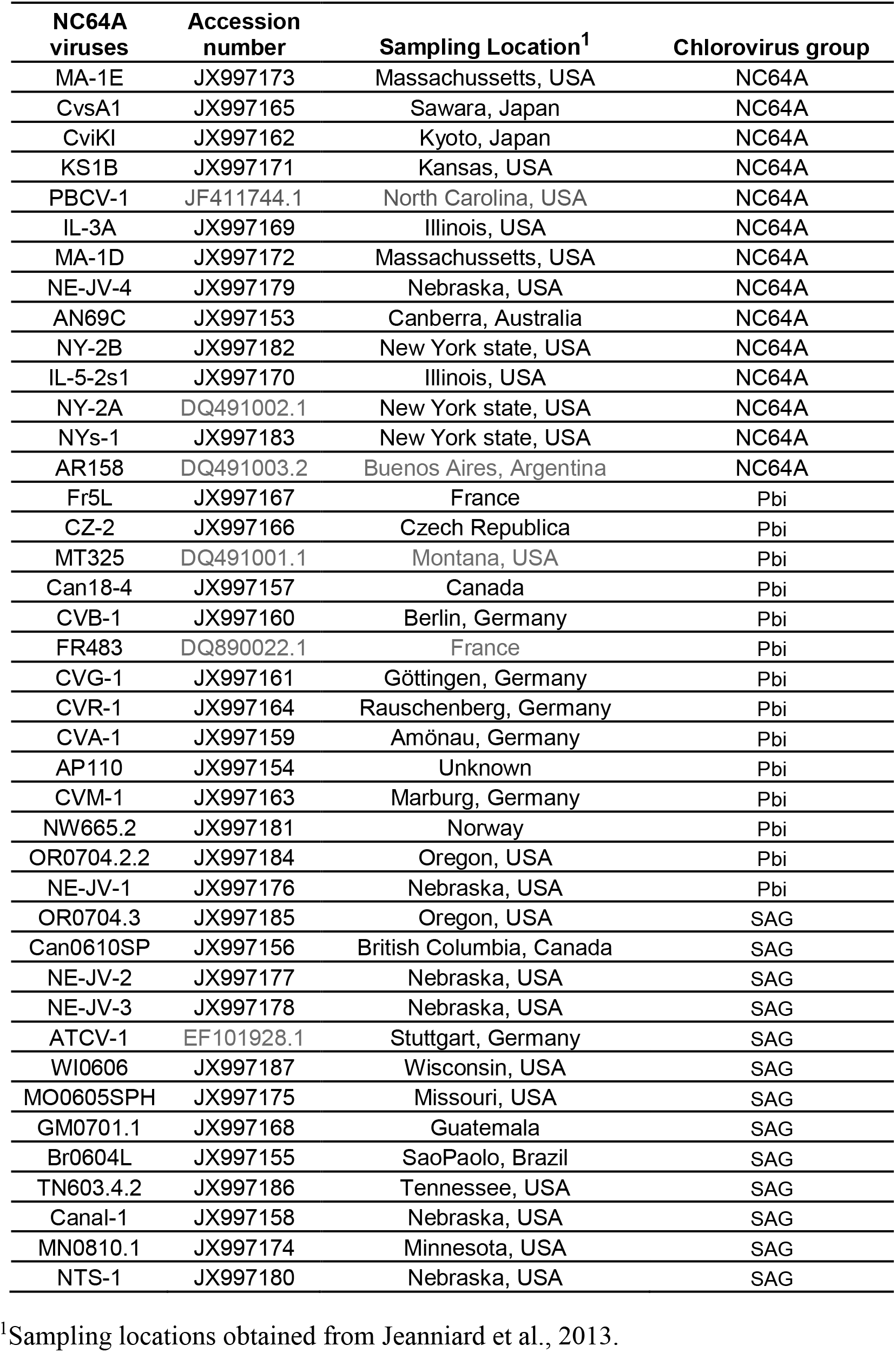
Accession numbers for 41 chloroviruses grouped into three clades that have three different hosts.

**Table S2.**
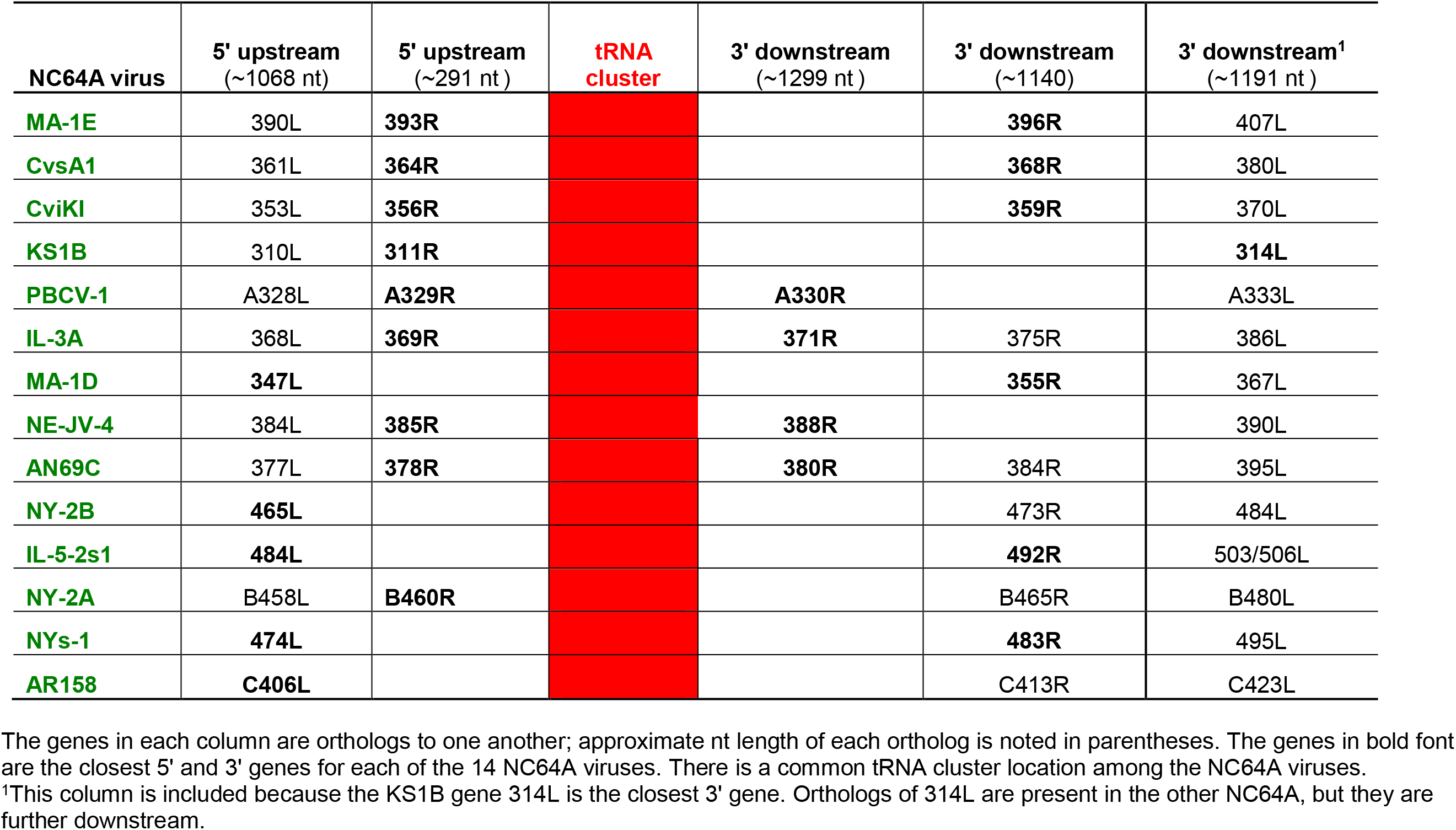
**A)** NC64A viruses: 5’ and 3’ genes closest to the tRNA gene cluster. **B**) SAG viruses: 5’ and 3’ genes closest to the tRNA gene cluster. **C**) Pbi viruses: 5’ and 3’ genes closest to the tRNA gene cluster.

**Table. S2B.**
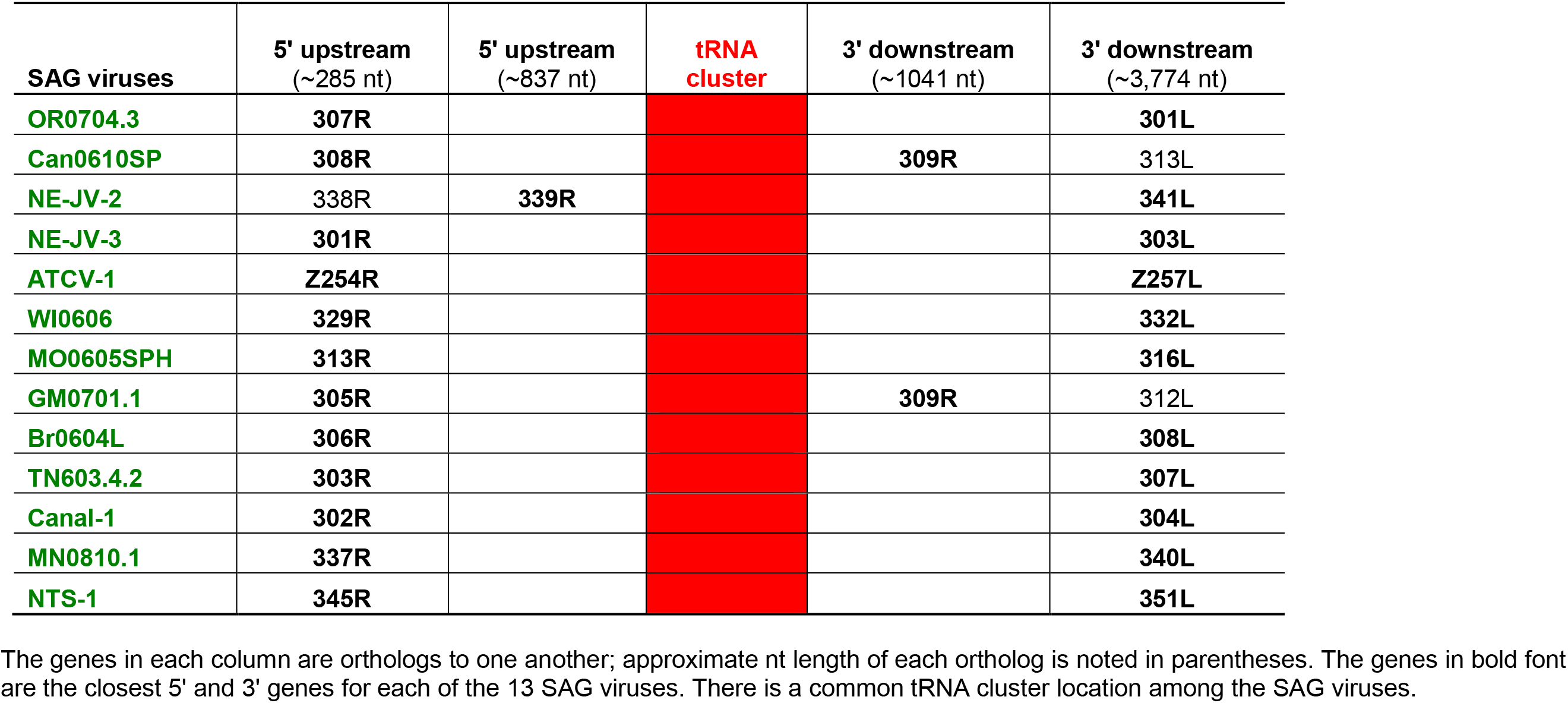
SAG viruses: 5’ and 3’ genes closest to the tRNA gene cluster.

**Table. S2C.** Pbi viruses: 5’ and 3’ genes closest to the tRNA gene cluster.

| Pbi viruses | 5' upstream<br>(~726 nt) | 5' upstream<br>(~270 nt) | 5' upstream<br>(870 nt) | 5' upstream<br>(267 nt) | tRNA<br>cluster | 3' downstream<br>(789 nt) | 3' downstream<br>(~297 nt)<sup>1</sup> | 3' downstream<br>(~420 nt) | 3' downstream<br>(~552 nt) |
| --- | --- | --- | --- | --- | --- | --- | --- | --- | --- |
| Fr5L | 393R | **397R** |  |  |  |  |  | **401L** |  |
| CZ-2 | 352R | **355R** |  |  |  |  |  | **358L** |  |
| MT325 | **M3422R** |  |  |  |  |  | **M344L** |  |  |
| Can18-4 | **414R** |  |  |  | 418R |  | **419L** | 422L |  |
| CVB-1 | **406R** |  |  |  | 408R |  | **411L** | 413L |  |
| FR483 | **N351R** |  |  |  |  |  | **N345L** | n356L |  |
| CVG-1 | **385R** |  |  |  |  |  | **390L** | 392L |  |
| CVR-1 | **400R** |  |  |  |  |  | **404L** | 406L |  |
| CVA-1 | **392R** |  |  |  |  |  | **396L** | 398L |  |
| AP110A | **403R** |  |  |  |  | **407R** | 411L | 413L |  |
| CVM-1 | **421R** |  |  |  |  |  | **425L** | 428L |  |
| NW665.2 | **375R** |  |  |  | 378R |  | **381L** | 383L |  |
| OR0704.2.2 | 349R | 352R | **353R** |  |  |  |  | **356L** |  |
| NE-JV-1 |  |  |  | **683R** | 688R |  |  |  | **690L** |

The genes in each column are orthologs to one another; approximate nt length of each ortholog is noted in parentheses. The genes in bold font are the closest 5' and 3' genes for each of the 13 SAG viruses. There is a common tRNA cluster location among the Pbi viruses, with the exception of NE-JV-1, which continues to be unique. Several viruses have protein-encoding genes within the tRNA cluster, which are not orthologous to one another. The HGT events that led to these gene insertions (blue) resulted in the loss of one or more tRNA genes in each case.
<sup>1</sup>MT325 and FR483 have longer genes (525 nt) but the core of 297 is present with high identity.

**Figure. S1A.**
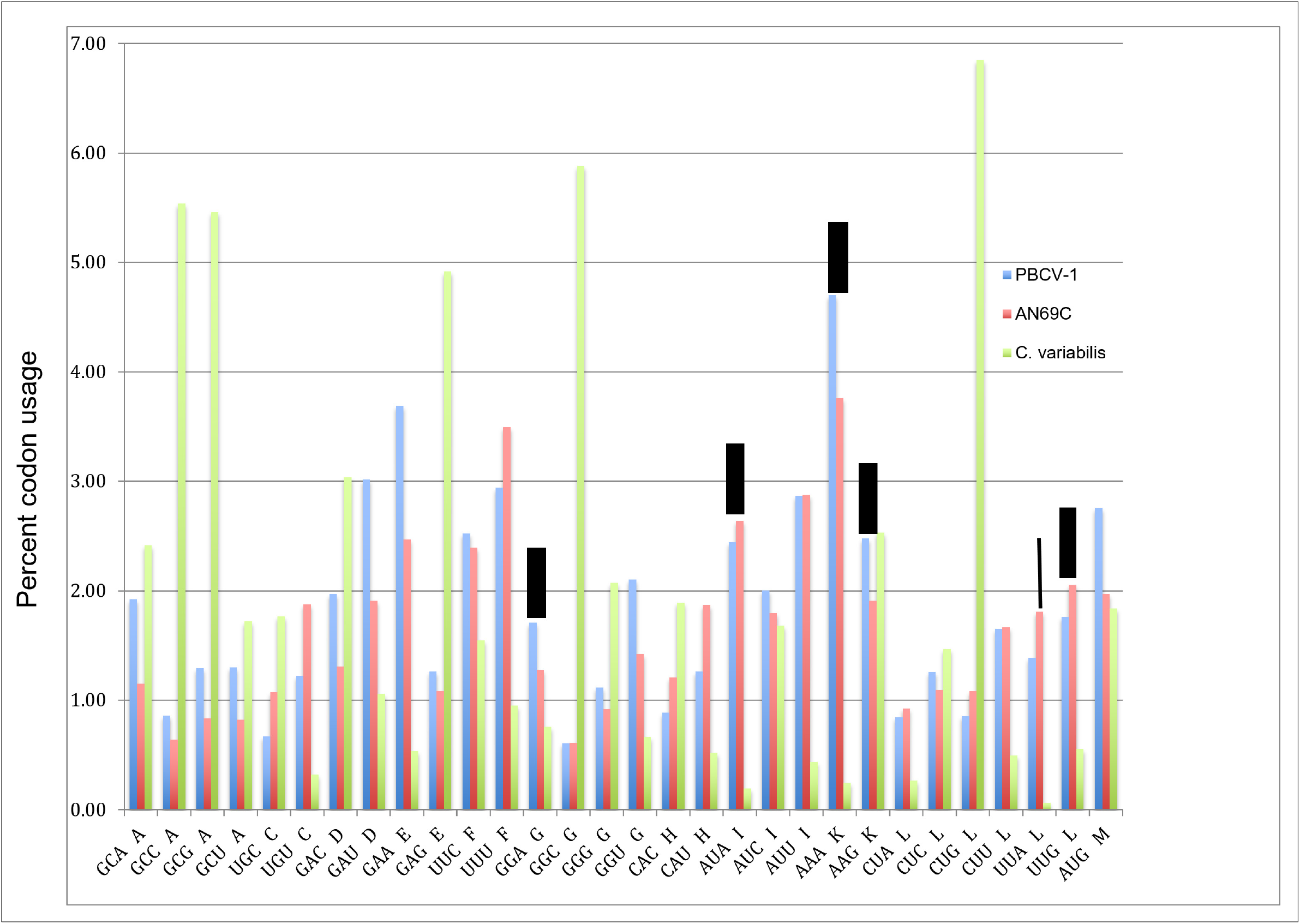
**A**) First half of genetic code book, comparing frequency of codon usage in the NC64A viruses PBCV-1 and AN69C to their host, *C. variabilis* NC64A. **B**) Second half of genetic code book, comparing frequency of codon usage in the NC64A viruses PBCV-1 and AN69C to their host, *C. variabilis* NC64A. Wide bars denote the codons recognized by tRNAs encoded by PBCV-1 and AN69C. Narrow dashed bar denotes codon recognized by tRNA encoded by PBCV-1 but not AN69C.

**Figure. S1B.**
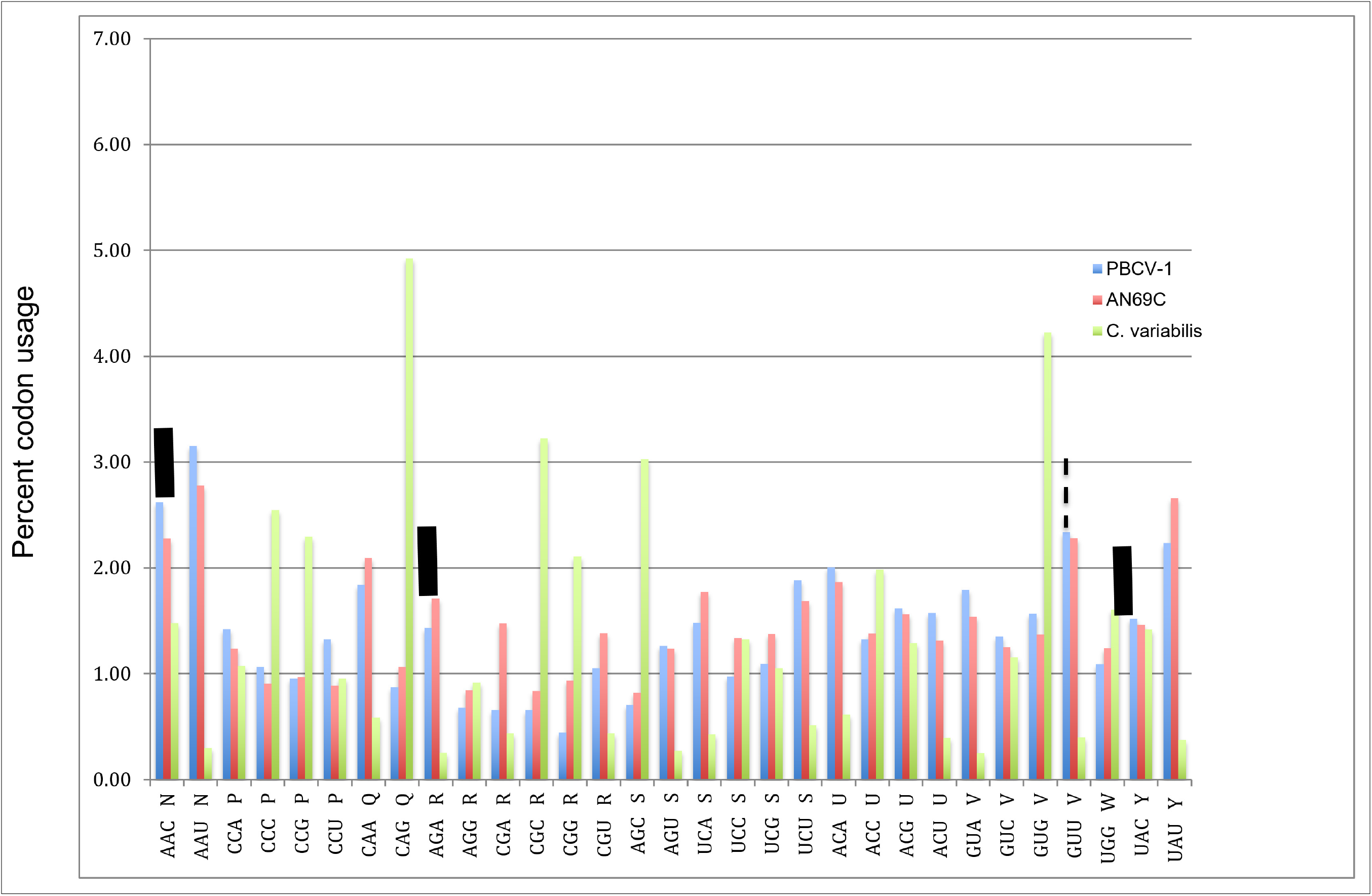
Second half of genetic code book, comparing frequency of codon usage in the NC64A viruses PBCV-1 and AN69C to their host, *C. variabilis* NC64A. Wide bars denote the codons recognized by tRNAs encoded by PBCV-1 and AN69C. Narrow dashed bar denotes codon recognized by tRNA encoded by PBCV-1 but not AN69C.

